# Investigation of Soft Magnetic Material Cores in Transcranial Magnetic Stimulation Coils and the Effect of Changing Core Shapes on the Induced Electric Field in Small Animals

**DOI:** 10.1101/2022.12.08.519188

**Authors:** Mohannad Tashli, George Weistroffer, Aryan Mhaskar, Deepak Kumbhare, Mark S. Baron, Ravi L. Hadimani

## Abstract

Transcranial magnetic stimulation (TMS) is a safe, effective and non-invasive treatment for several psychiatric and neurological disorders. Lately, there has been a surge in research utilizing this novel technology in treating other neurological and psychiatric ailments. The application of TMS on several neurological disorders requires the induced electric and magnetic fields to be focused and targeted to a small region in the brain. TMS of a focal cortical territory will ensure modulation of specific brain circuitry without affecting unwanted surrounding regions. This can be achieved by altering the properties of the magnetic core material used for the TMS system. In this study, soft ferromagnetic materials having high permeability, high saturation magnetization and low coercivity have been investigated as TMS coil cores in finite element simulations. Also, magnetic field measurements have been carried out using different cores in the TMS coil.

Finite element analysis of the rat head model is carried out using Sim4life software while investigating variations associated with changing the ferromagnetic core material and shape in the coil. Materials proposed for the analysis in this study include Iron Cobalt Vanadium alloy (Fe-Co-V) also known as Permendur, Carbon Steel (AISI 1010) and Manganese Zinc ferrites (MnZn ferrites). Simulation results indicated significant magnetic field distribution variation when introducing a ferromagnetic core in TMS coil, concentrating the magnetic field to the targeted region in the rat head model without stimulating adjacent regions. It was observed that the v-tip sharpened core attained the highest magnetic field and best focality among other cores in simulations and experimentally.

## Introduction

Transcranial Magnetic Stimulation (TMS) is a safe, effective and non-invasive therapy for a few mental and psychiatric disorders [1] [2]. TMS is FDA approved treatment and is commonly applied to patients who do not respond to medications for the treatment of clinical depression, smoking cessation and obsessive-compulsive [3]–[7]. The development of electromagnetic neuromodulation approaches aimed at improving the efficacy of TMS devices for the treatment of mental illnesses has grown recently. For efficacious clinical outcomes, the electromagnetic pulses must target specific regions of the brain with a precise magnitude and distribution to prevent undesired brain stimulation and overstimulation during TMS treatment [8].

The most prominent challenge in TMS is achieving a certain magnetic flux density and electric field strength at a certain depth in the brain. It is known that magnetic flux density and electric field strength are maximal at the surface and attenuate with depth [9]–[11]. Most commercial TMS coils have not included cores in their coil designs, for example, Magstim, Nextim, Magventure and Brainsway coils [12]–[15] which will make it hard to use these coils on small animals, as these coils stimulate the whole rat brain rather than stimulating a specific region. There are few reports in the literature on the development of focused TMS coils for rats [16]. Some designs used angle-tuned coils to enhance focality and penetration depth [17]. However, there are no reports that have compared different ferromagnetic core materials and shapes to be used in TMS coils.

In our study, we focused on investigating varying core designs for TMS coils by changing core material and shapes to increase the stimulation strength and to be able to focus the magnetic field to a 1 mm^2^ area. Core design in TMS coils requires deliberate considerations in terms of permeability, saturation magnetization and coercivity. Magnetic loss within the coil core in the form of eddy currents, static hysteresis, and anomalous loss [18], due to the high fundamental frequency of TMS is also another consideration for TMS core materials. Three prospective materials were considered as a coil core in TMS application: Carbon Steel AISI-1010, Iron-Cobalt-Vanadium alloy and Manganese Zinc ferrite (MnZn). Magnetic properties of potential TMS core materials were investigated using a vibrating-sample magnetometer (VSM). All these chosen materials exhibit good magnetic properties in terms of high magnetic saturation, high relative permeability and low core losses.

Core shape in TMS coils plays an important role in magnetic field distribution and magnitude in the rat brain. We investigated four different core shapes in our simulations. Sim4life, a finite element analysis software was used to test the magnetic field variation on a rat head model using four different proposed core geometries; cylindrical, v-tip, inverted-tip and reduced diameter flat tip cores. A uniaxial hall probe was used to measure the magnetic field on the cylindrical and v-tip ferromagnetic cores. These cores have been chosen out of the four proposed shapes based on the simulation results, aiming to compare the best focality core against the lowest focality one.

## Methodology

We developed an anatomically accurate rat head model as reported in a previous publication [19] to be used in our simulations. TMS stimulation parameters in Sim4life simulations were set to match the parameters reported in Selvaraj *et al*. [20] in terms of the coil shape and stimulator strength. The stimulator and focused coil reported in Selvaraj *et al*. [20] were used to measure the generated magnetic field at different depths corresponding to the coil bottom surface using two types of cores.

### A. Rat Head Model

To develop anatomically accurate rat head model, magnetic resonance images (MRIs) and computed tomography (CT) images have been collected for the rat’s brain from the NITRC public database [21]. These MRI’s went through segmentation in Statistical Parametric Mapping (SPM) using the SIGMA brain atlases. Then Meshmixer and Meshlab programs have been used to correct any model flaws and decrease the file size [19]. Anatomically accurate rat head model is shown in Fig. 1. This model simulates different electrical conductivity, permittivity, and density for the rat’s skin, skull, cerebrospinal fluid (CSF) grey matter and white matter.

**Fig. 1.**
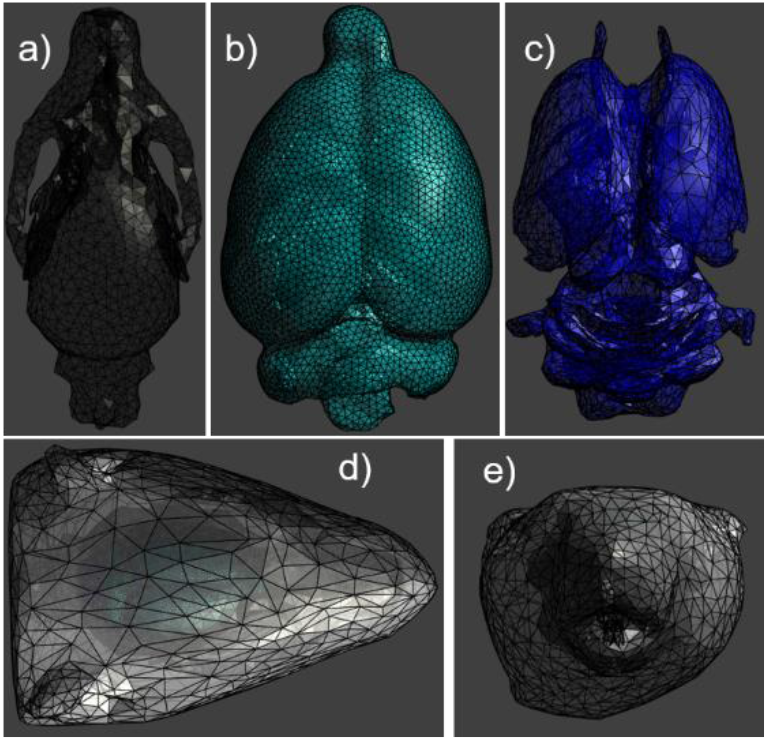
Anatomically accurate rat head model brain segments. a) skull, b) gray matter, c) white matter, d) skin and tissues and e) is the front view of the head model.

### B. Material characterization

The magnetic moment of different materials was measured using the Quantum Design Physical Property Measurement System (PPMS) vibrating sample magnetometer (VSM) option. The magnetic moment is measured as a function of magnetic field with a fixed temperature of 300K. Measurements have been performed on Carbon Steel AISI-1010, Iron-Cobalt-Vanadium alloy and Manganese Zinc ferrite MnZn with sample masses of 0.1383, 0.0223 and 0.0186 grams, respectively. We have applied a magnetic field ranging between −30,000 Oe to 30,000 Oe (−3T to 3T) on the Iron-Cobalt-Vanadium alloy and Carbon Steel samples while we applied a magnetic field ranging between −5,000 Oe to 5,000 Oe (−0.5T to 0.5T) on the MnZn ferrite sample. Then, the data is used to plot the M-H and B-H curves of the materials after calculating the magnetic flux density B (in Tesla) using equation 1

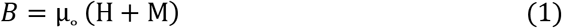

Where μ_o_ = 4π× 10^−7^ (vacuum permeability), H is the magnetic field in A/m and M is magnetization in A/m.

### C. Core Shapes

Four proposed core geometries have been investigated to maximize the magnetic field generated by the TMS coil taking into consideration the focality of these cores. The studied cores were inverted-tip (A), cylindrical (B), flat-tip with reduced diameter (C), and v-tip (D) (Fig. 2). All cores were 150 mm long with the largest diameter of 25.4 mm.

**Fig. 2.**
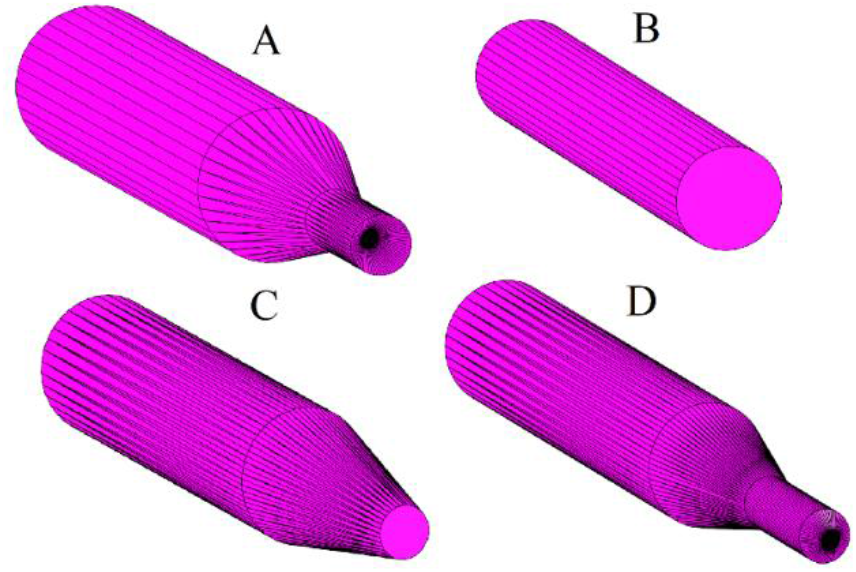
Proposed core geometries for TMS coils. A) Inverted tip core. B) Cylindrical core. C) Reduced diameter flat tip core. D) V-tip core

### D. Simulation

Sim4Life finite element analysis software (Zurich Med Tech, v6.2.1.4972) was used to solve for the magnetic H-field (A/m) from peak intensity stimulation at different depths of rat grey matter tissue from horizontal plane slices. The simulated coil and simulation parameters matched the work reported in Selvaraj *et al*. [20]. However, in our simulations, we used varying TMS stimulator core geometries shown in Fig. 2 to concentrate magnetic flux lines towards the center of the targeted stimulation area of the rat head.

In Sim4Life, the TMS stimulator with modified core and wound coil was positioned directly above the targeted rat head region (center of the grey matter of the brain) perpendicular to the horizontal plane. A visualization of the simulation set up for each coil configuration can be seen in Fig. 3. Material properties of the individualized segments and surrounding air were selected from the IT’IS LF database (IT’IS Foundation, v4.0) [22] for skin, skull, grey matter, white matter, cerebrospinal fluid (shown in Fig. 4). A static vector-potential simulation was executed with the amplitude of TMS set to 1000 amps to determine the magnetic H-field. This step was needed in order to account for varying magnetic permeability in the stimulator core. These simulation results were then used as a source to execute a magneto quasi-static simulation at 2.5 kHz [23]. Using a surface viewer of Sim4Life, the magnetic field output from the quasi-static simulation on the grey matter was observed from a top view and visualized.

**Fig. 3.**
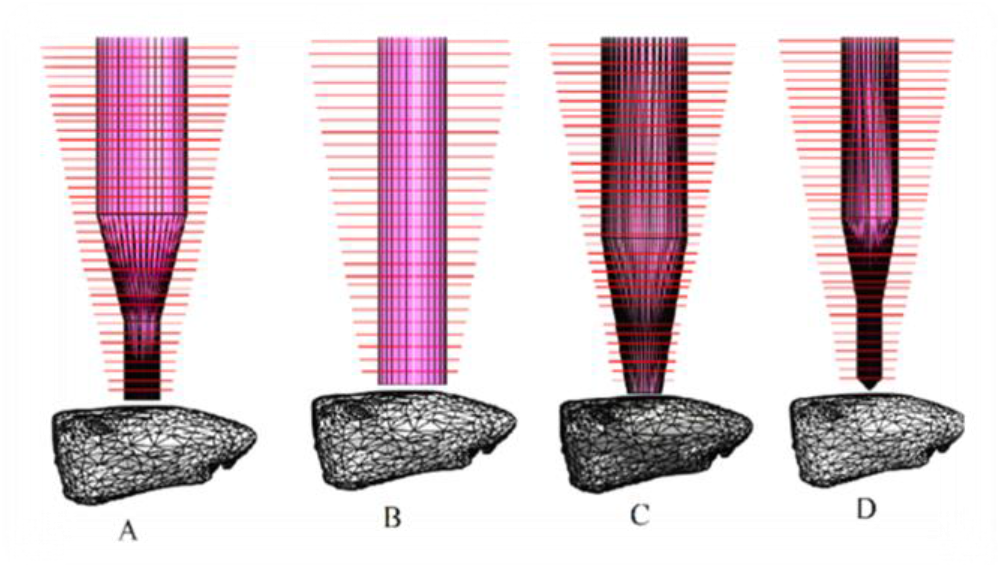
Simulation configuration for the 4 different core shapes in the TMS focused coil. A) Inverted tip core. B) Cylindrical core. C) Reduced diameter flat tip core. D) V-tip core.

**Fig. 4.**
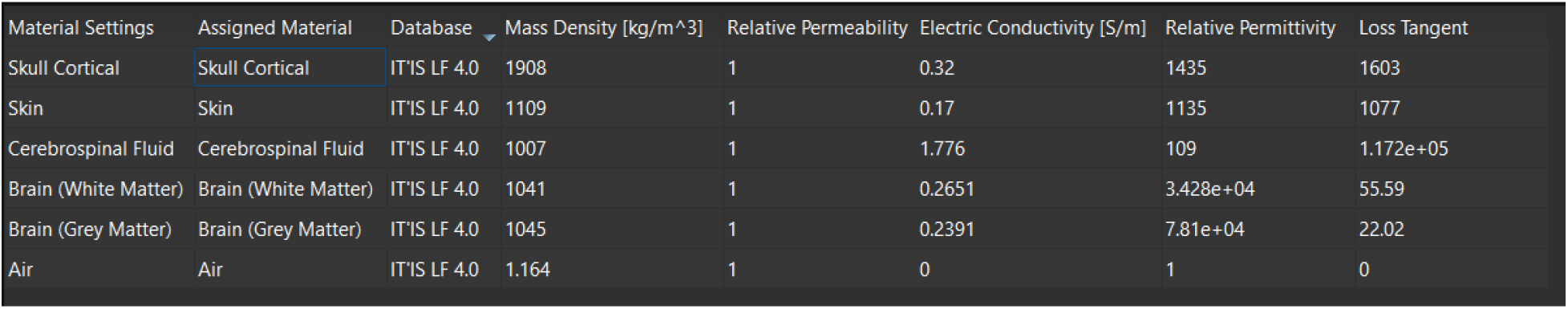
Rat’s brain layers material properties assigned in Sim4life simulations.

### E. Hall probe magnetic field measurements

A uniaxial hall probe is used to measure the magnetic field generated in the TMS focused coil. The TMS stimulator and focused coil used in this study were designed and published in a previous paper by Selvaraj [20]. The coil has 40 turns with a conical shape. The inductance of the coil with the v-tip carbon steel core is measured (65μH). The original circuit was designed to handle a current of 1000 Amps. We were able to increase that current to 1300 (Amps.) by changing the capacitor to a bigger one. Based on the simulation results, we have measured the magnetic field using the hall probe in the cylindrical and the v-tip ferromagnetic cores. The focused coil with ferromagnetic core magnetic field measurement setup are shown in Fig. 5.

**Fig. 5.**
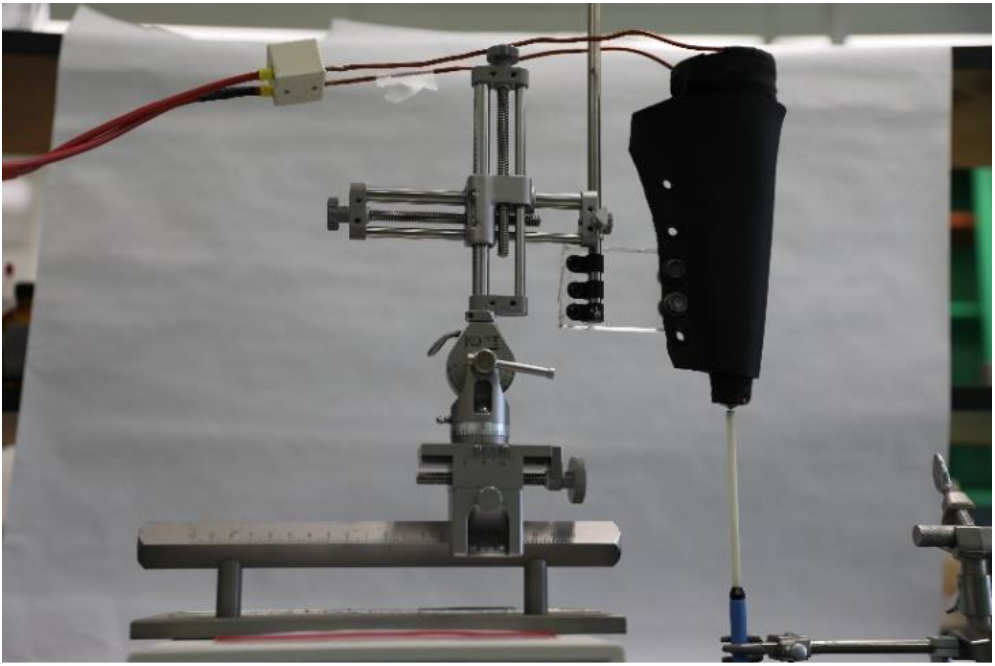
The TMS focused coil with a ferromagnetic core in magnetic field measurement setup using a hall probe.

## Results

### A. VSM

The B-H curve are plotted for the three materials in the study using the data generated after measuring magnetization vs. magnetic field in a VSM for the materials are shown in Fig. (6–8) and the M-H curves are shown in Fig. 9, Fig. 10 and Fig. 11. The measured saturation flux densities were 2.35 Tesla for the Iron-Cobalt-Vanadium alloy (Fig. 6) and 0.45 Tesla for the Manganese Zinc ferrite (Fig. 7) compared to 2.4 Tesla and 0.47 Tesla in their vendor data sheets [24], [25]. The saturation flux density for the Carbon Steel AISI-1010 was 2 Tesla (Fig. 8) which was the same in SysLibrary of ANSYS. The reported relative permeability of materials of interest were 3350 for the Iron-Cobalt-Vanadium alloy [25], 2000 for the MnZn ferrite [24] and 667 for the Carbon Steel AISI-1010 in SysLibrary of ANSYS.

**Fig. 6.**
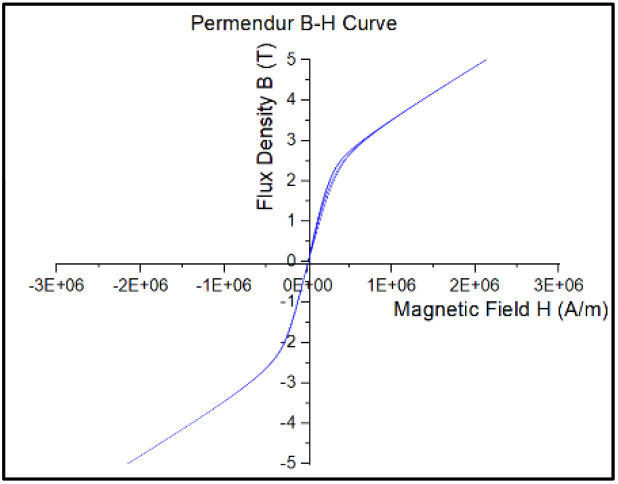
Iron-Cobalt-Vanadium alloy (Fe-Co-V alloy) B-H curve.

**Fig. 7.**
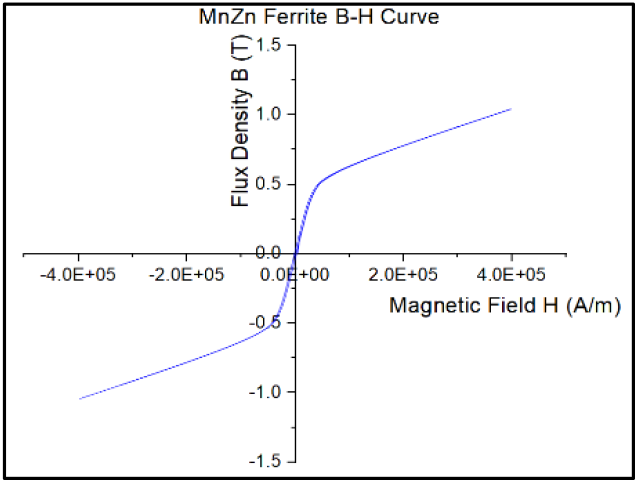
Manganese Zinc ferrite (MnZn ferrite) B-H curve.

**Fig. 8.**
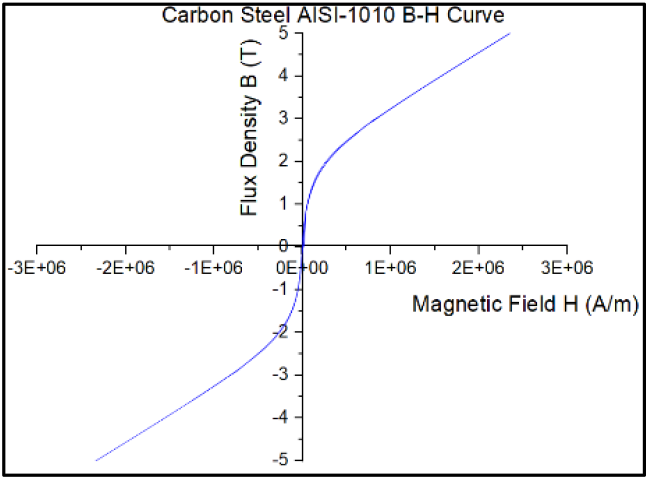
Carbon Steel AISI-1010 B-H curve.

**Fig. 9.**
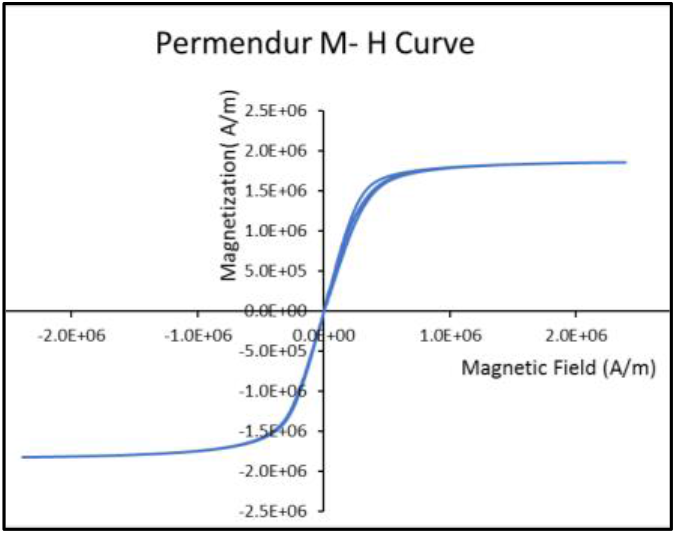
Iron-Cobalt-Vanadium alloy (Fe-Co-V alloy) M-H curve

**Fig. 10.**
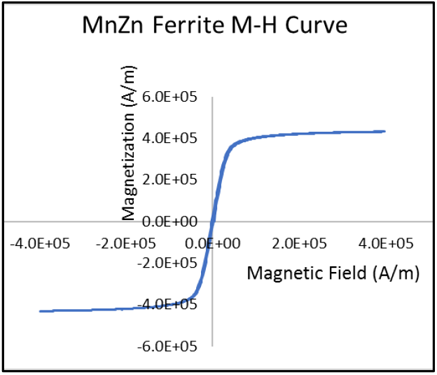
Manganese Zinc ferrite (MnZn ferrite) M-H curve

**Fig. 11.**
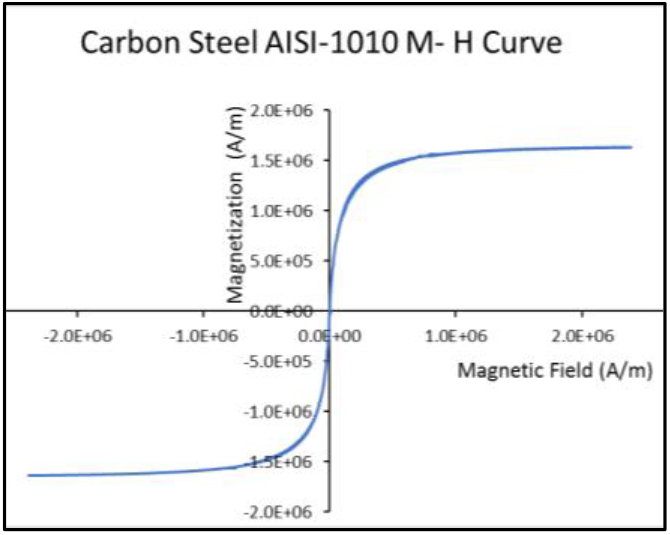
Carbon Steel AISI-1010 M-H curve.

### B. Sim4life Simulations

Sim4life simulations for the TMS-focused coil with the four ferromagnetic core shapes have been performed using one of the three ferromagnetic materials, the carbon steel AISI-1010 with a relative permeability μr = 667. The results of the Sim4Life simulations showed the greatest focused magnetic H-field strength using v-tip core geometry, peaking at a strength of 2.6 MA/m at the site of stimulation directly underneath where the coil was positioned. Additionally, this core configuration had the least amount of extraneous gray matter stimulated, showing high focality at the desired site of stimulation. The visualization of the magnetic H-field in A/m on the rat gray matter for all core shapes can be seen in Fig. 12. The corresponding coil shape letter in Fig. 3 is used to signify which results image corresponds to which core shape. The first row of the simulated head models in Fig.11 shows the results when the H-field scale is fixed for all 4 simulations in the range of 0 to 2.6×10^6^A/m indicating 0 with black and 2.6×10^6^A/m with white color scale. In the second row of results, the H-field distribution was normalized for each simulation corresponding to their maximum value to demonstrate the focality for each core.

**Fig. 12.**
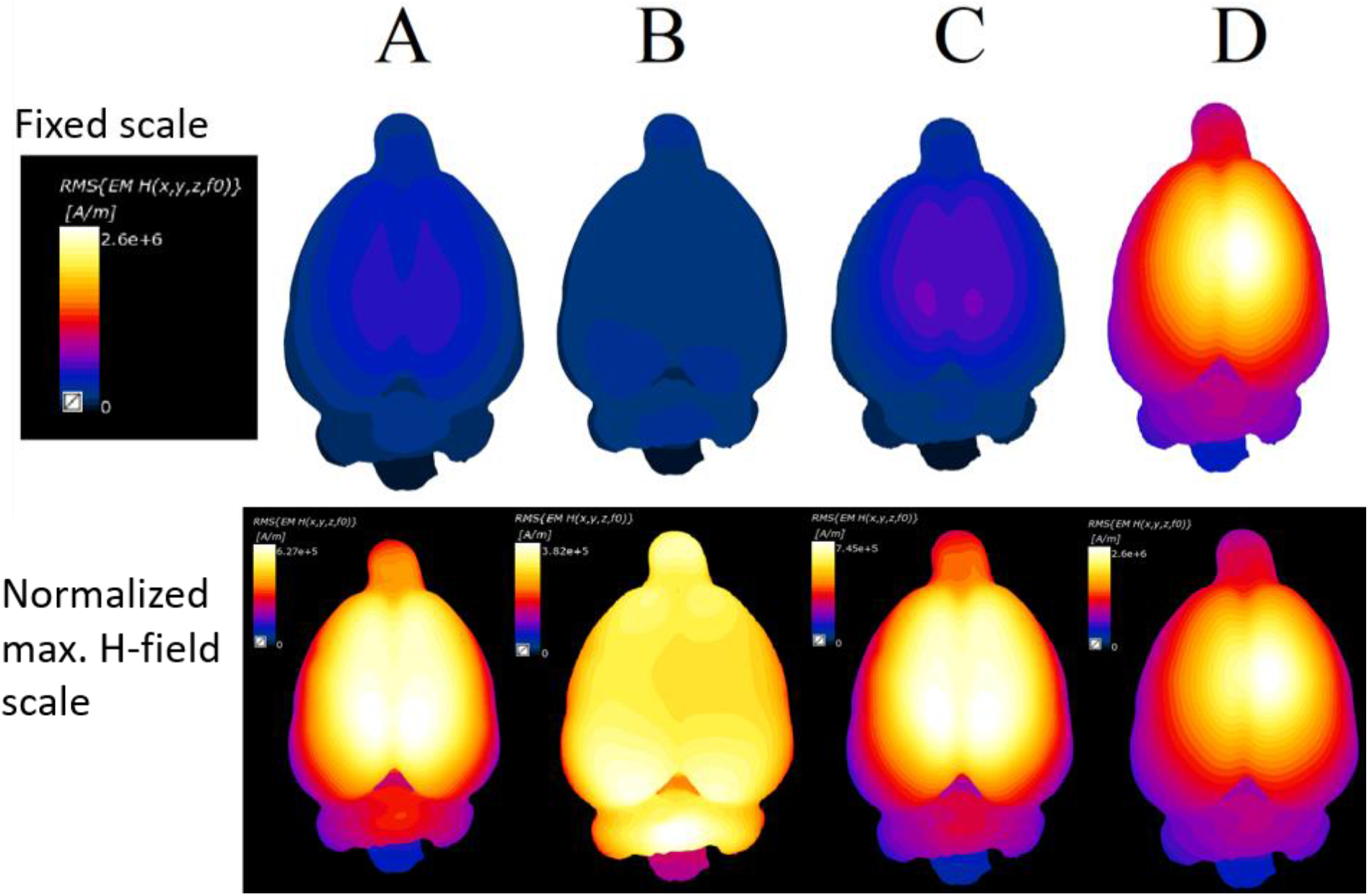
Simulation results showing H-field strength distribution on the rat’s gray matter using a fixed H-field scale (first row). Normalized H-field results to the maximum value for each core simulation (second row). Letters above the simulated head models represent the core shape used in that simulation. Inverted-tip (A), cylindrical (B), flat-tip with reduced diameter (C), and v-tip (D)

For a better view of the focality for the 4 cores, spatial lines of the magnetic field have been plotted at the surface of the rat head model’s grey matter for the different core geometries as shown in Fig. 13. The magnetic field for each core has been normalized to better assess the focality for each core. The v-tip core indicated the narrowest width among other cores which demonstrates its focality. The full-width-half-maximum of the induced magnetic fields have been calculated for the 4 different cores. The FWHM values were 32.9, 24.8, 21.6 and 14 mm for the cylindrical core, inverted-tip core, flat-tip core and the v-tip core, respectively.

**Fig. 13.**
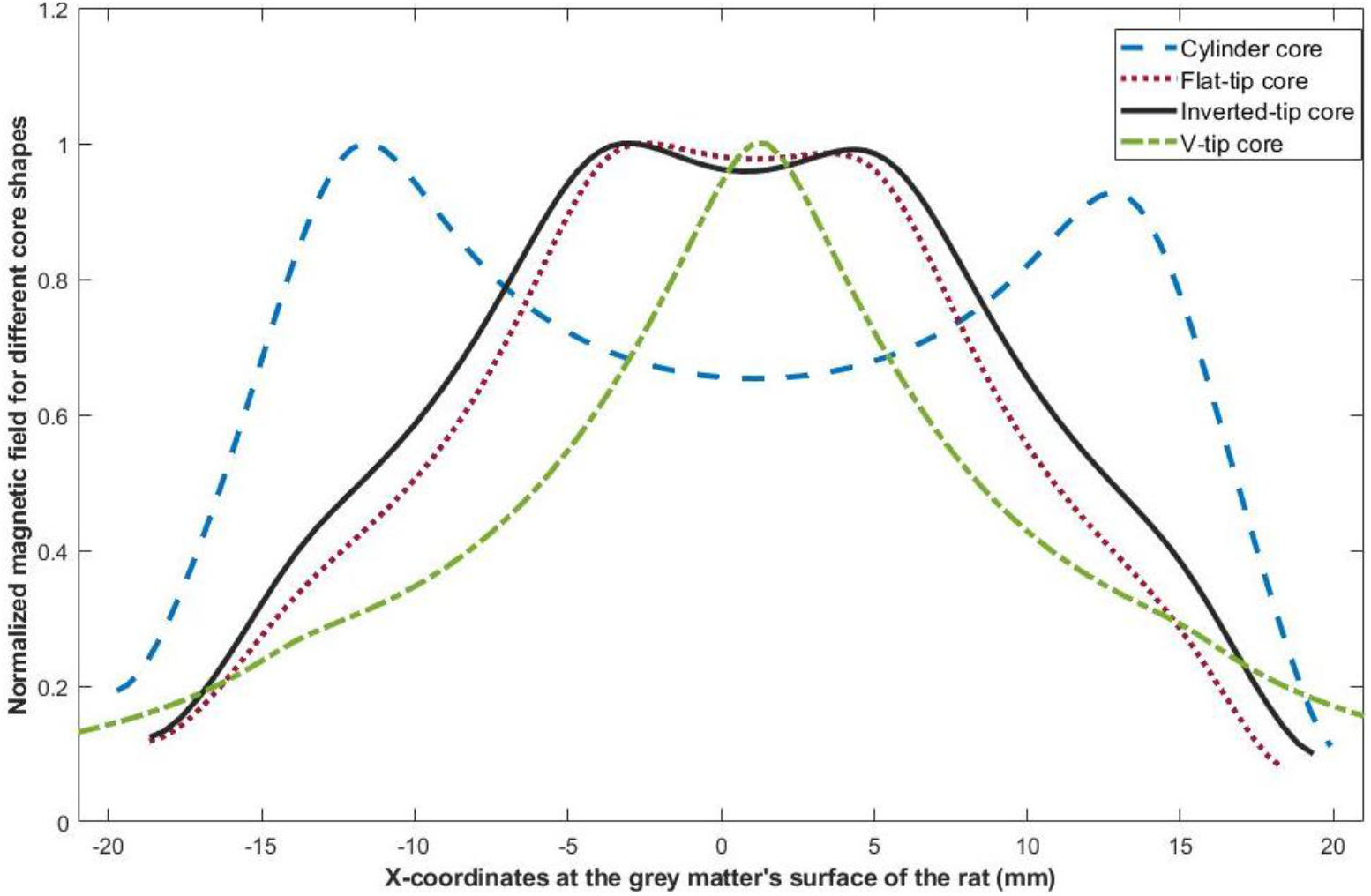
Normalized magnetic field for the 4 different core geometries to demonstrate the focality for each core.

### C. Hall probe magnetic field measurements

Inserting a carbon steel core in the TMS coil improved the measured magnetic field significantly using the hall probe. The magnetic field measurements right below the coil are listed in Table 1 comparing results using the two cores chosen (the v-tip and cylindrical ferromagnetic cores). The v-tip sharpened coil gave us the highest magnetic field measurement right below the coil. At 2 mm depth below the coil, the measured magnetic field dropped in magnitude in both situations; using the cylindrical and v-tip cores as shown in Table 2. The magnetic field drop at the center of the v-tip core was significantly higher than the drop using the cylindrical core as it dropped from 7.72E+05 A/m to 2.92E+05 A/m. However, the v-tip core measured magnetic field was still the highest. Additionally, the rise time of the pulse was 90 μs considering 1000 (Amps.) Flowing through the coils.

**Table 1.**
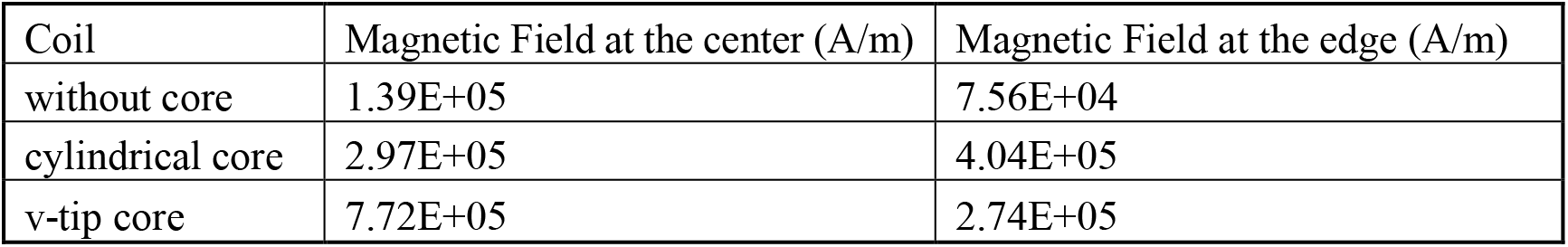
Magnetic field measurements right below the coil surface with and without core.

**Table 2.** Magnetic field measurements at 2 mm distance below the coil surface with two types of cores.

| Coil | Magnetic Field at the center (A/m) | Magnetic Field at the edge (A/m) |
| --- | --- | --- |
| cylindrical core | $2.51\text{E}+05$ | $2.86\text{E}+05$ |
| v-tip core | $2.92\text{E}+05$ | $2.13\text{E}+05$ |

## Discussion

In this study, we investigated the effect of using different ferromagnetic cores in TMS coils in terms of core shape and material. Three prospective ferromagnetic materials of interest have been chosen; Iron-Cobalt-Vanadium alloy, Carbon steel (AISI-1010) and MnZn ferrite.

As for the core shape selection, the study evaluated through Sim4life simulations four possible core shapes (inverted-tip, cylindrical, flat-tip with reduced diameter, and v-tip). Based on simulations results, the v-tip and cylindrical ferromagnetic cores were actually used to perform the magnetic field measurements using the hall probe, as the v-tip core achieved the best magnetic field focality and magnitude, while the cylindrical core had the lowest focality.

Magnetic measurements of these cores showed excellent magnetic properties, high relative permeability, high saturation and low coercivity. Of the three materials mentioned above, AISI 1010 was shortlisted to characterize the tip shapes using magnetic field measurements using the hall probe. Magnetic measurements using the hall probe for the other two materials will be conducted in the future.

It has been noticed that the measured point’s density in the linear part of the B-H curves was low. To measure the slope of the linear region and calculate the initial relative permeability of the material, more data are required in the linear part of the curve when we first magnetize the material. Moreover, the demagnetization factor was not taken into consideration for the measured samples, and that’s due to the irregularity in the geometries of the samples, which will affect the relative permeability significantly [26]–[28]. Hence, our calculated relative permeabilities calculated from the VSM measurements (Fig. 6–8) were lower than that in the data sheets. The magnetic fields in simulations appeared to be higher than that in the experiment; that’s due to some restrictions in Sim4life simulation software. In simulations, we assigned the maximum relative permeability for the core material which is a constant value while in reality, the relative permeability changes as the slope of the B-H curve change with increasing the magnetic field. Moreover, the electrical conductivity of the core material cannot be changed in Sim4life, and that will cause the Eddy-current losses in the core minimal compared to the experimental work.

## Conclusion

Finite element analysis of the rat head model is carried out using Sim4life software to investigate variations associated with changing the coil core material and geometry. Simulation results showed that introducing a ferromagnetic core in the TMS-focused coil improved the focality of that coil significantly. The v-tip ferromagnetic core achieved the highest magnetic field and best focality compared to other cores.

Magnetic field measurements have been performed on the Carbon Steel AISI-1010 cylindrical and v-tip cores in the TMS-focused coil using a hall probe. The v-tip core still achieved the highest magnetic field at the bottom surface of the coil, as well as at 2 mm distance below the coil. We believe that using Iron-Cobalt-Vanadium alloy as a v-tip core in the TMS-focused coil will enhance the coil performance significantly, for its high saturation flux density and relatively high relative permeability.

V-tip cores increase focality in TMS coils, especially for small animals. This TMS core coil will enable stimulating small regions in the brains of small animals for testing neuronal circuits. Thus, coils with cores can help develop new TMS procedures in treating psychological and psychiatric disorders by targeting extremely small regions otherwise hard to target with commercial TMS coils.

## Acknowledgments

Authors acknowledge Commonwealth Cyber Initiative Central Virginia Node Grant # FP00010500. This study was also partially funded by the Merit Review Award, U.S. Department of Veterans Affairs to Dr. Mark Baron as a principal investigator. Grant number 2I01BX001147-05A2. Authors would like to thank Dr. Ivan Carmona from Northwestern University for assisting in core shapes design and in the stimulator modification.

